# Long-term potentiation prevents ketamine-induced aberrant neurophysiological dynamics in the hippocampus-prefrontal cortex pathway *in vivo*

**DOI:** 10.1101/763540

**Authors:** Cleiton Lopes-Aguiar, Rafael N. Ruggiero, Matheus T. Rossignoli, Ingrid de Miranda Esteves, José Eduardo Peixoto Santos, Rodrigo N. Romcy-Pereira, João P. Leite

**Author notes:** these authors contributed equally to this work (Cleiton Lopes-Aguiar, Rafael N. Ruggiero).

## Abstract

N-methyl-D-aspartate receptor (NMDAr) antagonists such as ketamine (KET) produce psychotic-like behavior in both humans and animal models. NMDAr hypofunction affects normal oscillatory dynamics and synaptic plasticity in key brain regions related with schizophrenia, particularly in the hippocampus and the prefrontal cortex. In contrast, long-term potentiation (LTP) induction is known to increase glutamatergic transmission. Thus, we hypothesized that LTP could mitigate the electrophysiological changes promoted by KET. We recorded HPC-PFC local field potentials and evoked responses in urethane anesthetized rats, before and after KET administration, preceded or not by LTP induction. Our results show that KET promotes an aberrant delta-high-gamma crossfrequency coupling in the PFC and an enhancement in HPC-PFC evoked responses. LTP induction prior to KET attenuates changes in synaptic efficiency and prevents the increase in cortical gamma amplitude comodulation. These findings are consistent with evidence that increased efficiency of glutamatergic receptors attenuates cognitive impairment in animal models of psychosis. Therefore, high-frequency stimulation in HPC may be a useful tool to better understand how to prevent NMDAr hypofunction effects on synaptic plasticity and oscillatory coordination in cortico-limbic circuits.

## Introduction

The hippocampal-prefrontal cortex (HPC-PFC) monosynaptic pathway is implicated in cognitive functions, such as working memory, decision making, and spatial-temporal processing^1,2^. Dysfunctional connectivity within HPC-PFC circuits is associated with the pathophysiology and genetic predisposition of schizophrenia^3–5^. In schizophrenia, HPC-PFC connectivity is decreased during working memory tasks and increased in resting state^5–7^. Such effects may be mediated, at least in part, by N-methyl-D-aspartate receptor (NMDAr). In fact, NMDAr binding is reduced in schizophrenic patients in both HPC and PFC, and administration of an NMDAr antagonist, such as ketamine, can induce psychotic symptoms in healthy patients and an increase in resting state HPC-PFC connectivity^8–10^. This NMDAr hypofunction also affects synaptic plasticity, inducing impairments in critical circuits, such as HPC-PFC, promoting cognitive symptoms by pathologic neural activity^11–13^. However, the relationship between synaptic plasticity in HPC-PFC circuits and schizophrenia is not fully understood.

In rodents, ketamine and other NMDAr antagonists are widely used as a translatable pharmacological model capable of inducing psychotic-related behaviors, such as hyperlocomotion, working memory impairments, prepulse inhibition disruption, and abnormal social interaction^14,15^. Several neurophysiological features of the HPC-PFC pathway are associated with psychotic-like behaviors induced by NMDAr antagonists. *In vivo* experiments showed that ketamine increased gamma power in the PFC, affected the synaptic transmission efficiency in the HPC-PFC pathway, and disrupted long-term potentiation (LTP)^16–18^.

Interestingly, studies indicate that allosteric modulation of NMDAr or α-amino-3-hydroxy-5-methyl-4-isoxazolepropionic acid receptor (AMPAr) can prevent NMDAr antagonist-induced impairments^19–21^. The allosteric modulation of NMDAr reverts working memory deficit induced by ketamine and is capable of increasing NMDAr function in PFC^22,23^. Consistently, clozapine, an atypical antipsychotic, has been shown to increase NMDAr currents, both by inhibiting type-A glycine transporter and binding to glycine-site NMDAr^19,24^. Furthermore, clozapine is also capable of preventing ketamine impairment on LTP induction of the HPC-PFC pathway^18,25^. Regarding AMPAr, its positive allosteric modulation reversed impairments in HPC-PFC short-term synaptic plasticity and increase delta oscillation in the PFC^26^. Although the clinical antipsychotic effects of allosteric modulation in glutamatergic receptors are not conclusive^27,28^, it is known that induction of synaptic plasticity increases AMPAr availability and NMDAr efficiency, which could revert NMDAr antagonist impairments^29,30^. Assuming that (1) NMDAr antagonists affect synaptic plasticity^16–18,31–33^; (2) allosteric modulation of glutamatergic receptors prevents psychotic-like effects,^19–24^ and (3) induction of LTP modulates both AMPAr and NMDAr activity^34–36^; we hypothesized that a previous increase of glutamatergic efficiency by LTP induction can preclude NMDAr antagonist effects on HPC-PFC pathway.

To test our hypothesis, we explored the effects of acute ketamine injection on the spontaneous oscillatory and evoked activities in the HPC-PFC pathway in anesthetized animals, and then, we examined if a previous LTP induction could modulate the neurophysiological effects of ketamine on HPC-PFC interaction. We first, classified basal brain state oscillations and then compared PFC field responses induced by HPC stimulation and HPC-PFC oscillatory coupling before and after ketamine treatment. Then we tested the modulatory effect of deep brain stimulation (LTP at HPC-PFC synapses) on ketamine-driven oscillatory patterns. Our findings show that ketamine induces a state of increased gamma frequency power and abnormal cross-frequency coupling in the PFC, followed by an enhancement of HPC-PFC evoked responses. Although LTP induction also increases gamma frequency activity, it prevented ketamine-induced aberrant oscillatory coupling and potentiation of HPC-PFC synaptic transmission. Our data suggests that modulation of synaptic plasticity in HPC-PFC circuits might be a useful tool to better understand how to prevent dysfunctions induced by NMDAr blockade.

## Results

To investigate ketamine effects on the field activity of the HPC-PFC pathway we first characterized the brain oscillatory activity during urethane anesthesia. In figure 1 we show distinct alternating oscillatory patterns elicited by urethane anesthesia: 1) deactivated states (DEA), characterized by high amplitudes and slow rhythms in the LFP resembling Slow-Wave sleep patterns (Figure 1C); and 2) activated states (ACT), described by an increase of faster frequency bands power (>4 Hz), which resembles REM oscillatory activity ^37^ (Figure 1C). To separate brain states, we analyzed the RMS and zero-crossings value of each epoch (Figure 1D). In figure 1E we represent the spectra content of each brain state. Consistent with previous reports^37,38^, we show that in DEA there is a predominance of delta oscillations (1 Hz), while in ACT epochs there is a significant decrease in delta (PFC: t_(7)_=9.712,*p*<0.0001; HPC: Wilcoxon matched-pairs signed rank test, n=8,*p*=0.0078), and an increase in theta (PFC: t_(7)_=13.15, *p*<0.0001; HPC: t_(7)_=4.474, *p*=0.0029), low-gamma (PFC: t_(7)_=4.054, *p*=0.0048; HPC: t_(7)_=2.357,*p*=0.0506) and high-gamma (PFC: t_(7)_=4.964,*p*=0,0016; HPC: Wilcoxon matched-pairs signed rank test, n=8, *p*=0.1094) relative power for both PFC and HPC.

**Figure 1.**
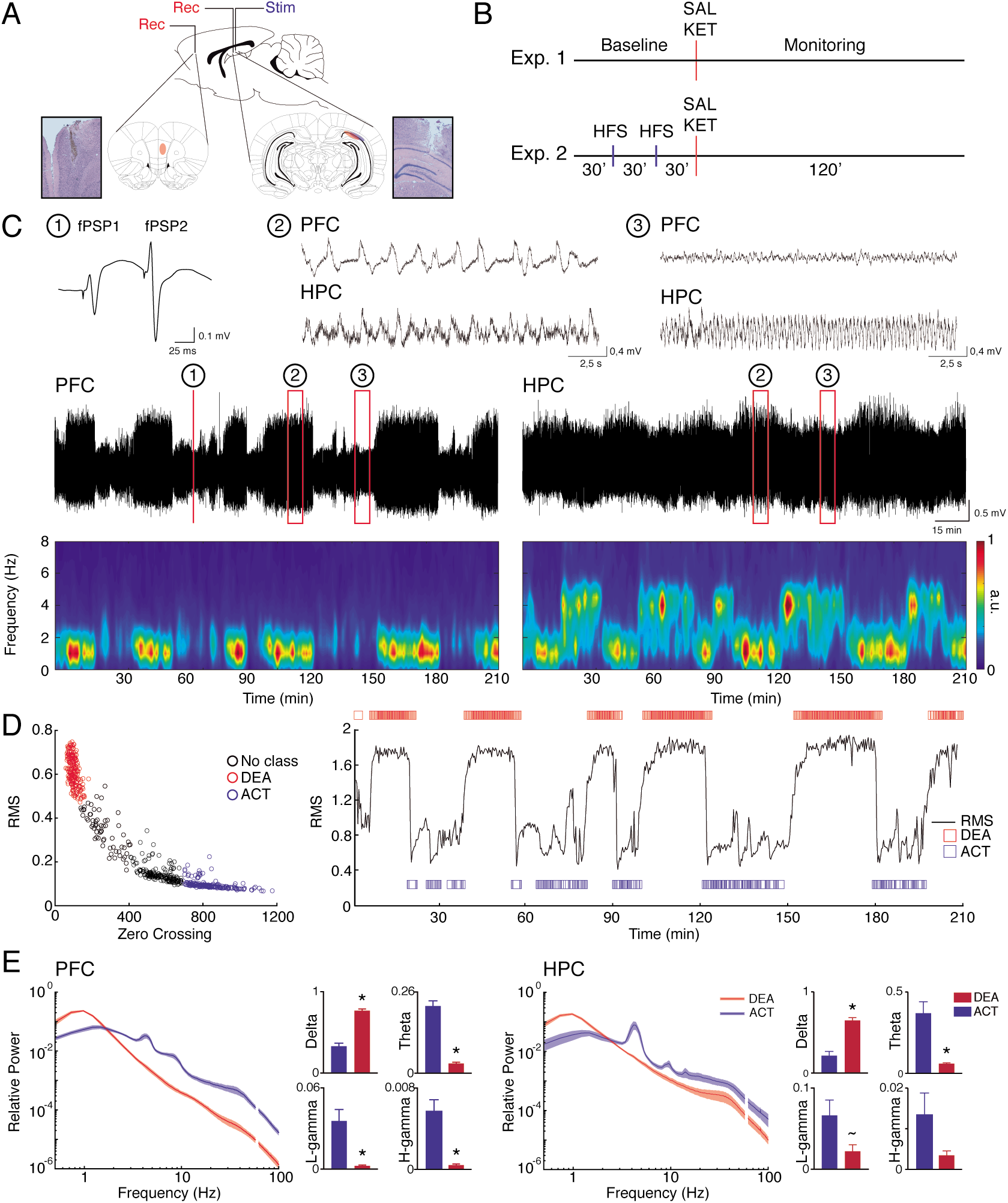
Oscillatory states under urethane anesthesia. (A) Positioning of recording and stimulus electrodes and representative electrolytic lesions in Nissl-stained coronal sections. (B) Experimental design. (C) Representative evoked fPSPs in the PFC (1), and raw LFP during deactivated (2) and activated (3) states in PFC (top) and HPC (bottom). Representative 210 min recording of a raw LFP (middle) and spectrograms (bottom) in the PFC and HPC showing spontaneous brain state alternation during urethane anesthesia. (D) Brain state classification. Scatter plots of RMS values and the number of zero crossing were used to classify brain states in deactivated (DEA) and activated (ACT) epochs (left). RMS and state classification in a representative recording (right). (E) Relative power spectral densities in the PFC and HPC (line plots) during different brain states (SAL, n=8). In both regions, activated states present theta oscillation and an increase in gamma frequencies, while in deactivated state there is a predominance of delta oscillation (bar plots). *p<0.05.

### HPC-PFC connectivity under urethane anesthesia

In figure 2A we measured synchrony between HPC and PFC through spectral coherence. We show that while DEA epochs show synchrony in delta rhythms, spectral coherence during ACT state demonstrates a peak in theta oscillations (4 Hz, Figure 2A). These patterns of synchronicity closely follow alternation of brain states (Figure 2A). We further explored state connectivity by performing cross-correlation and Granger causality analyses. In Figure 2B we show that the peak of crosscorrelation in delta frequency (predominant oscillation on DEA epochs) have a positive lag (2 ms), while theta oscillations indicate a negative lag (−22 ms). Granger analysis reveal that ACT periods show theta oscillations with the HPC leading the PFC (t_(7)_=2.4909, *p*=0.0415), while on DEA epochs we found a peak in delta oscillations (~1.5 Hz) with the PFC driving the HPC oscillations (t_(7)_=7.7401, *p*=0.0001). These results suggest that during ACT states hippocampal theta coordinates activity in the PFC, while in DEA periods delta oscillations from the PFC coordinates HPC slow activity. We also explored phase-amplitude coupling on each state. We found a prominent coupling of the delta phase with high-gamma amplitude during DEA oscillatory activity in the PFC (Figure 2C). This coupling was significantly higher than a uniform empirical distribution (shuffled surrogate data) and the one present in ACT epochs (Kruskal-Wallis test: H_(3)_=16.81, *p*=0.0002, Dunn’s post-hoc test *p*=0.0002 for DEA vs. ACT and p=0.0157 DEA vs. Surrogate data; Figure 2C). Comodulation maps did not reveal a clear cross-frequency coupling (CFC) during ACT states (data not shown).

**Figure 2.**
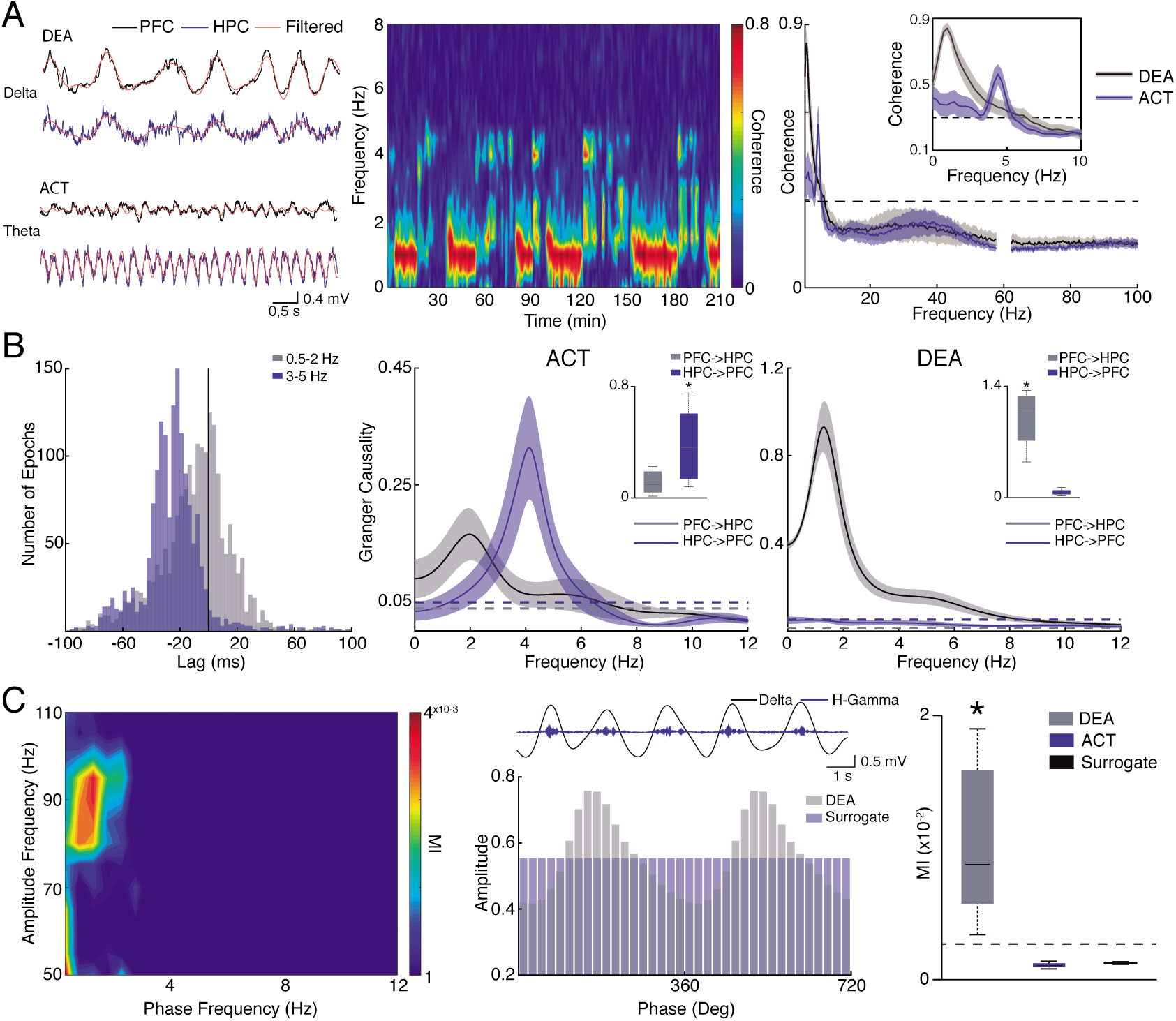
Brain state modulation of HPC-PFC connectivity. (A) Raw and filtered LFP during DEA (delta) and ACT (theta) in the PFC and HPC epochs (left). Representative coherogram for the entire recording (middle) showing delta (~1 Hz) and theta (~4 Hz) synchrony alternation during brain states. Averaged spectral coherence plot (right). Inset showing coherence peak ~4 Hz and ~1 Hz during ACT and DEA epochs, respectively. The dashed line represents 95% confidence interval of bootstrapped data. (B) Histogram of cross-correlogram lags (peak) of delta and theta oscillations in DEA an ACT (left). Granger causality in the ACT (middle) and DEA states (left). The dashed line represents 95% confidence interval of permuted data. Insets: box plots distribution of maximum Granger theta (left) and delta (right) power, showing significant causality in the HPC-PFC direction during activated state (left), and PFC-HPC direction during deactivated state. (C) Representative comodulation map showing delta-high-gamma coupling during deactivated state (left). Middle: Representative filtered LFP (top) and mean high-gamma amplitude as a function of delta phase in DEA state and surrogate data (bottom). Mean delta-high-gamma coupling strength (MI) during brain states (right) and surrogate data. MI is significantly increased during deactivated states. All data correspond to SAL group (n=8). *(p<0.05).

### NMDAr blockade effects on HPC-PFC connectivity

In figure 3B we illustrate the ketamine effects on field potential of the HPC and PFC. We show that NMDAr blockade leads to an increase in evoked field post-synaptic potential (fPSP) amplitude (F_(20,260)_=1.611 significant interaction between groups *p*=0.050; Bonferroni post-hoc test are bottom of the graphs) and a robust reduction of paired-pulse facilitation (PPF) (F_(20,260)_=8.205 significant interaction between groups *p* =<0,001). Figure 3C shows a representative raw LFP from both regions indicating ketamine effects on brain oscillations. As described in other studies^17,33,39^, we verified that NMDAr blockade increased low-gamma and high-gamma power in the PFC independent of brain state (DEA: low-gamma: t_(6)_=2.327, *p*=0.0589; high-gamma: t_(6)_=3.423, *p*=0.0141; ACT: low-gamma: t_(5)_=4.108, *p*=0.0093; high-gamma: t_(5)_=3.258, *p*=0.0225; Figure 3D). In PFC, no significant effects were observed in other frequency bands, while in HPC there is a slightly increase in gamma power (Figure S1A). Interestingly, systemic ketamine appears to increase the probability of DEA state. In figure 3E we demonstrate the probability density function of DEA epoch occurrence in all animals in the KET group compared with animals of the SAL group. While the probability of SAL group oscillates between low and high values during the recording, this oscillation pattern is reduced after ketamine injection, with the probability of a DEA epoch occurrence kept high for at least 30 min after systemic injection (see also Figure S2). This increase in DEA oscillatory activity is not accompanied by alterations in synchrony between the HPC-PFC LFP accounting for epoch separation. As we show in figure 3F there are no significant changes between spectral coherence of DEA or ACT states comparing before or after ketamine injection. However, as shown in a representative coherogram, systemic ketamine leads to a persistent synchronization in delta, which follows the increase in DEA epochs.

**Figure 3.**
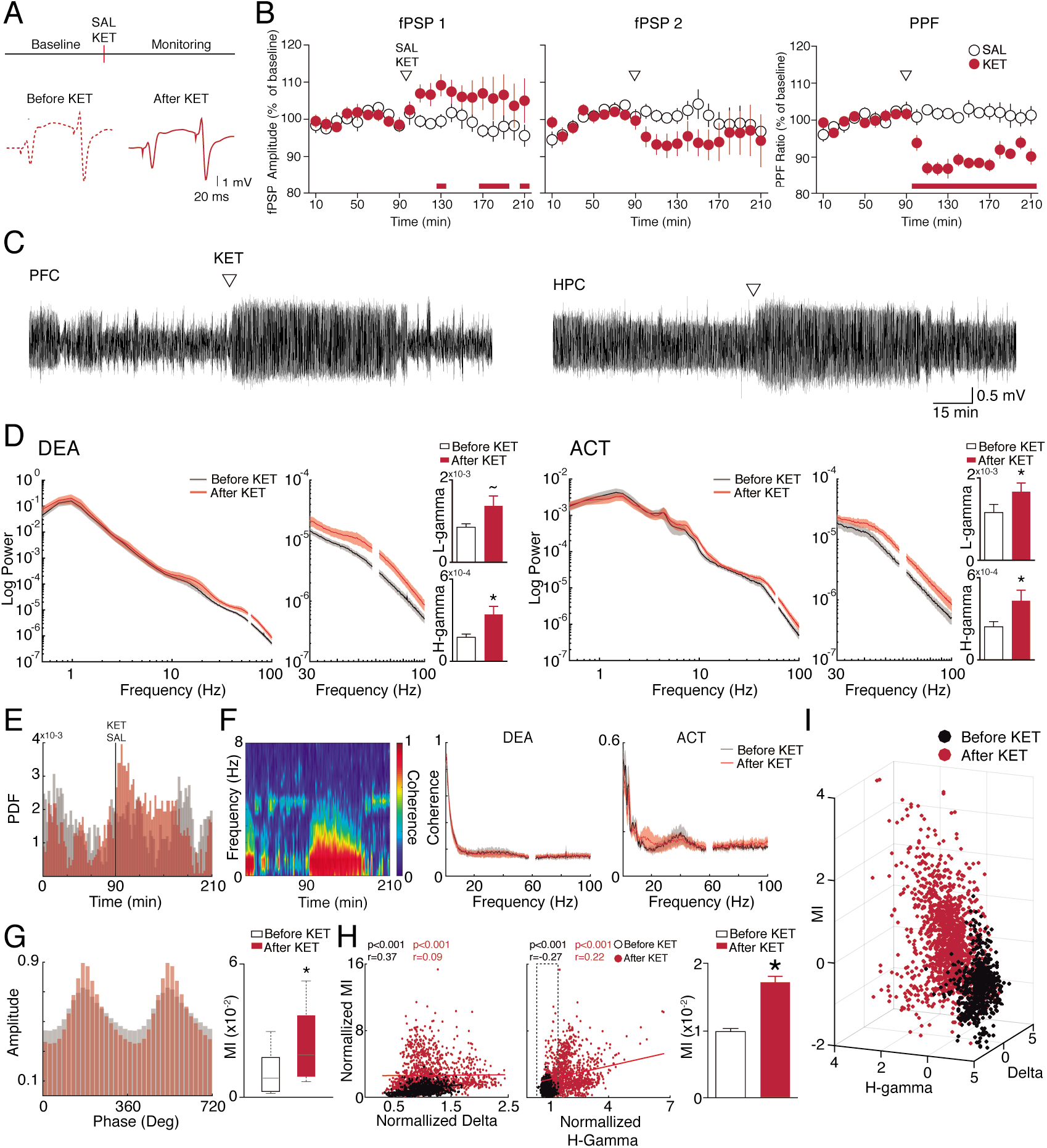
Brain state-dependent modulation of KET effects on HPC-PFC connectivity. (A) Experimental design for SAL (n=8) and KET (n=7) groups (top). Representative fPSPs of baseline (left) and after Ketamine injection (right). (B) Effects of ketamine injection on fPSPs (top and middle) and paired-pulse ratio (bottom) shown in 10 min blocks as mean ± standard errors. Data are presented as ratios from the baseline mean amplitude. Red horizontal bars on the bottom of graphs indicate post-hoc differences (p<0.05). (C) Representative raw in a ketamine injected rat. (D) Averaged PSDs in the PFC before and after ketamine injection in DEA (left) and ACT (right) states. There is a significant increase in low and high-gamma power after ketamine injection (bar plots). (E) Probability density function of DEA epochs along the recording in SAL and KET groups. (F) Representative coherogram in a KET rat (left) showing the increase of deactivated epochs. Averaged spectral coherence plots in deactivated (middle) and activated (right) states reveals that ketamine does not affect HPC-PFC synchrony. (G) Mean high-gamma amplitude as a function of delta phase in SAL and KET group (left). MI is significantly increased in deactivated states after ketamine injection. (H) Correlation of baseline normalized delta (left) and high-gamma (middle) power and MI before and after ketamine injection. Bar plot (right) compare DEA MI before and after ketamine injection in epochs which high-gamma power did not increase compared to baseline, showing that CFC increases independent of changes in power. (I) Three-dimensional scatter plot of z-scored MI, delta and high-gamma power showing a clear distinction of epochs before and after ketamine injection. *(p<0.05), ~(p=0.0589).

### NMDAr blockade promotes a different oscillatory state in the PFC

We examined if ketamine could influence DEA phase-amplitude coupling in the PFC. In figure 3G we represent high-gamma amplitude as a function of delta phase comparing DEA epochs before and after ketamine injection. We show here that NMDAr blockade increases cross-frequency coupling between delta and high-gamma activity (t_(6)_=4.947, *p*=0.0026). To investigate if the increase of the CFC was dependent on power changes promoted by ketamine we performed a linear correlation between DEA baseline-normalized MI and delta or high-gamma power values. Figure 3H shows a weak correlation between delta or high-gamma with MI values after ketamine injection (*r* = 0.09 for delta and *r*=0.22 for high-gamma). To further investigate if increase in CFC could be influenced by high-gamma power after ketamine we compared MI averaged values before and after the drug. For this, we only used post-ketamine epochs with high-gamma within 95% confidence interval of the pre-ketamine high-gamma distribution. We show that even epochs with no increase in high-gamma values compared to baseline present an average significant enhancement in CFC (Figure 3H right; t_(882)_=8.0693, *p*<0.001), indicating that ketamine effects in delta-high-gamma coupling may be independent of power increase in these frequencies. We confirmed this result with bootstrap analysis using 1000 repetitions and controlling for the number of trials. These data indicate that ketamine promotes an oscillatory state that is different from the traditional DEA state in the PFC. Figure 3I illustrates all the 20s DEA epochs from the KET group in a 3D plot showing how ketamine modify brain oscillatory activity in DEA epochs increasing gamma frequency band and delta-high-gamma coupling.

### LTP increases gamma activity in the PFC

In figure 4 we show the effects of high-frequency stimulation (HFS) stimulation on field potential of the PFC. As shown previously by our group^40–42^, HFS protocol induces a stable LTP in the HPC-PFC pathway for at least 120 min (fPSP1; F_(20,240)_=13.761, *p*<0.0001). LTP effects were also seen in the amplitude of the fPSP2 (paired pulse stimulation; F_(20,240)_=11.470, *p*<0.0001). However, the increase in the fPSP2 is relatively lower than the one in fPSP1, which is seen as a reduction in the PPF ratio at least for 60 min after HFS (F_(20,240)_=5.621, *p*<0.0001; Figure 4B). Figure 4C demonstrates the effects of HFS on LFP of the PFC. LTP induction produced an increase in low and high-gamma on the PFC restricted to DEA epochs (low-gamma: Mann-Whitney test, U=30, *p*=0.0006; high-gamma: Mann-Whitney test, U=27, *p*=0.0003). No significant alterations were observed at gamma frequencies in the HPC of LTP groups (Figure S1B). Interestingly, the increase in high-gamma power was not related to an enhancement of delta-high-gamma coupling. As shown in figure 4D modulation index did not differ from before or after LTP induction (p>0.05).

**Figure 4.**
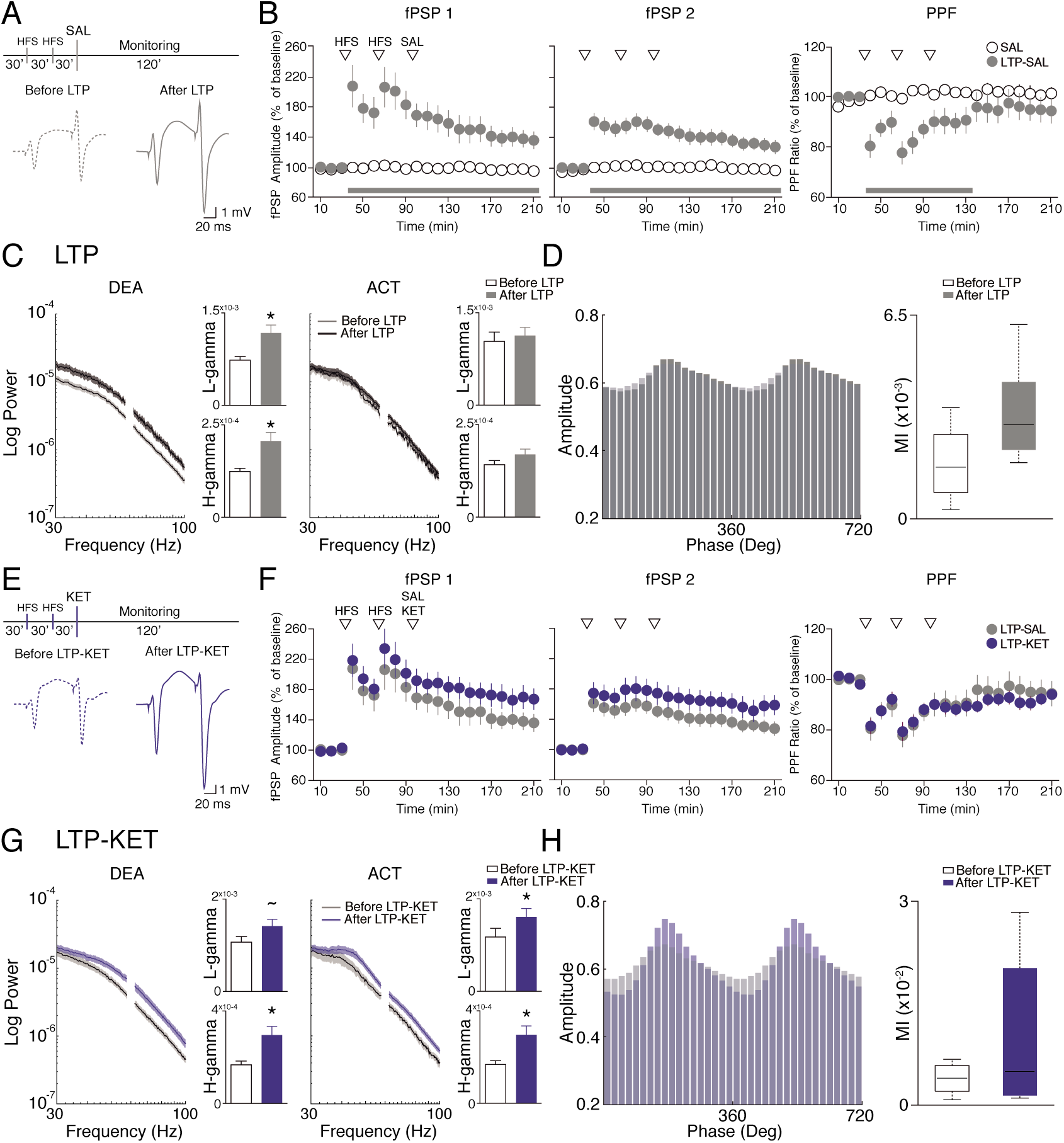
LTP modulation of Ketamine effects on HPC-PFC connectivity. Experimental design for LTP-SAL (A) and LTP-KET groups (E) (top) and representative fPSPs of baseline (left) and after HFS (right). (B) Effects of HFS (LTP-SAL group n=15) on fPSPs (left and middle) and paired-pulse ratio (right) shown in 10 min blocks as mean± standard errors. Data are presented as ratios from the baseline mean amplitude. (C) PSDs of gamma frequencies before and after HFS, in the DEA (left) and ACT states (right). Bar plots display mean and standard errors of low (top) and high-gamma power (bottom). There is a significant increase in gamma power after HFS in the DEA states. No differences were found in the activated state. (D) Mean high-gamma amplitude as a function of delta phase before and after HFS (left). MI in deactivated state is not affected by LTP induction (right). (F) LTP induction precludes the ketamine enhancement of fPSPs and PPF (LTP-KET, n=9). No statistical differences were observed between LTP and LTP-KET groups. (G) PSDs of gamma frequencies before and after Ketamine on LTP-KET group, in the deactivated (left) and activated states (right). There is a significant increase in high-gamma and a statistical trend in low-gamma power in the deactivated state (left). In the activated state there is significant increase in both low and high-gamma power (right). (H) Mean high-gamma amplitude as a function of delta phase before and after Ketamine in the LTP-KET group (left). MI in deactivated state is not affected in the LTP-KET group (right). *(p<0.05), ~(p=0.06).

### Ketamine effects are attenuated by prior LTP

To examine whether LTP was able to modulate the effects of ketamine upon PFC responses, we applied HFS prior to KET or vehicle administration (LTP-KET and LTP-SAL groups) and compared the results. Figure 4F indicates that ketamine treatment following LTP did not produce an increase in evoked potentials on the PFC either on fPSP1, fPSP2 or PPF comparing with control group (*p*>0.05). Similar to what we observed in the LTP-SAL group, LTP-KET also presented an increase in low and high-gamma activity in the DEA epochs of the PFC (low-gamma: t_(7)_=2.229, *p*=0.0611; high-gamma: t_(7)_=2.931, *p*=0.0220). While these effects are also seen in the LTP group LTP-KET show an increase in gamma frequency also during ACT states (low-gamma: t_(8)_=7.166, *p*<0.0001; high-gamma: t_(8)_=7.709, *p*<0.001; Figure 4G). These alterations appear to be related with the ketamine administration, since it was observed in the KET group as well (Figure 3D). Despite the high-gamma power increase, no significant alterations were observed in the delta-high-gamma coupling in DEA states (Figure 4H), indicating that LTP induction prevents ketamine effects on PFC CFC.

### LTP prevents ketamine-induced aberrant oscillatory brain state

We next compared the effects of ketamine followed or not by LTP. Figure 5A shows the absolute MI value across the entire recording in the KET and LTP-KET groups. We demonstrate that LTP induction attenuates ketamine effects on delta-high-gamma coupling especially in the first 30 minutes after drug injection. We next compared the baseline-normalized MI values for only DEA epochs in the initial 30 min after ketamine. Our data indicates that LTP attenuates the effects of ketamine on enhancing CFC coupling in the PFC (t_(12)_=2.589, *p*=0.0237). Figure 5B shows that KET and LTP-KET epochs after ketamine injection can be distinguished based on their electrophysiological features. We used principal component analysis for dimensionality reduction of electrophysiological features, and then applied a quadratic discriminant analysis to classify epochs in the two groups. For the quadratic function we used the first three principal components, which have an explained variance of 79.21% (Figure 5C). Our classification model with training data (filled circles) and then cross-validated with test data (open triangles) gave a classification accuracy of 85.56% accuracy. This data show that ketamine effects can be distinguished when preceded by LTP based on their electrophysiological characteristics. We next compared the epochs mean score from each group for the first three principal component (Figure 5D). Our results revealed that KET has a higher score on PC1 (*p*<0.0001), while LTP-KET has a higher score in PC2 (*p*=0.0069) and PC3 (*p*=0.0001). Interpreting the correlation coefficients (loadings) of the original variables with the first three principal components we observe that PC1 has a high correlation with CFC values (r=0.7696, *p*<0.0001), high-gamma (r=0.5975, *p*<0.0001) and delta activity (r=0.8963, *p*=0.0001). PC2 and PC3 have high correlation with theta (r=0.8368, *p*<0.0001) and high-gamma (r=0.6750, *p*<0.0001), respectively, and negative correlation with fPSP (r=- 0.5214, *p*<0.0001 and r=-0.5375, *p*<0.0001, respectively).

**Figure 5.**
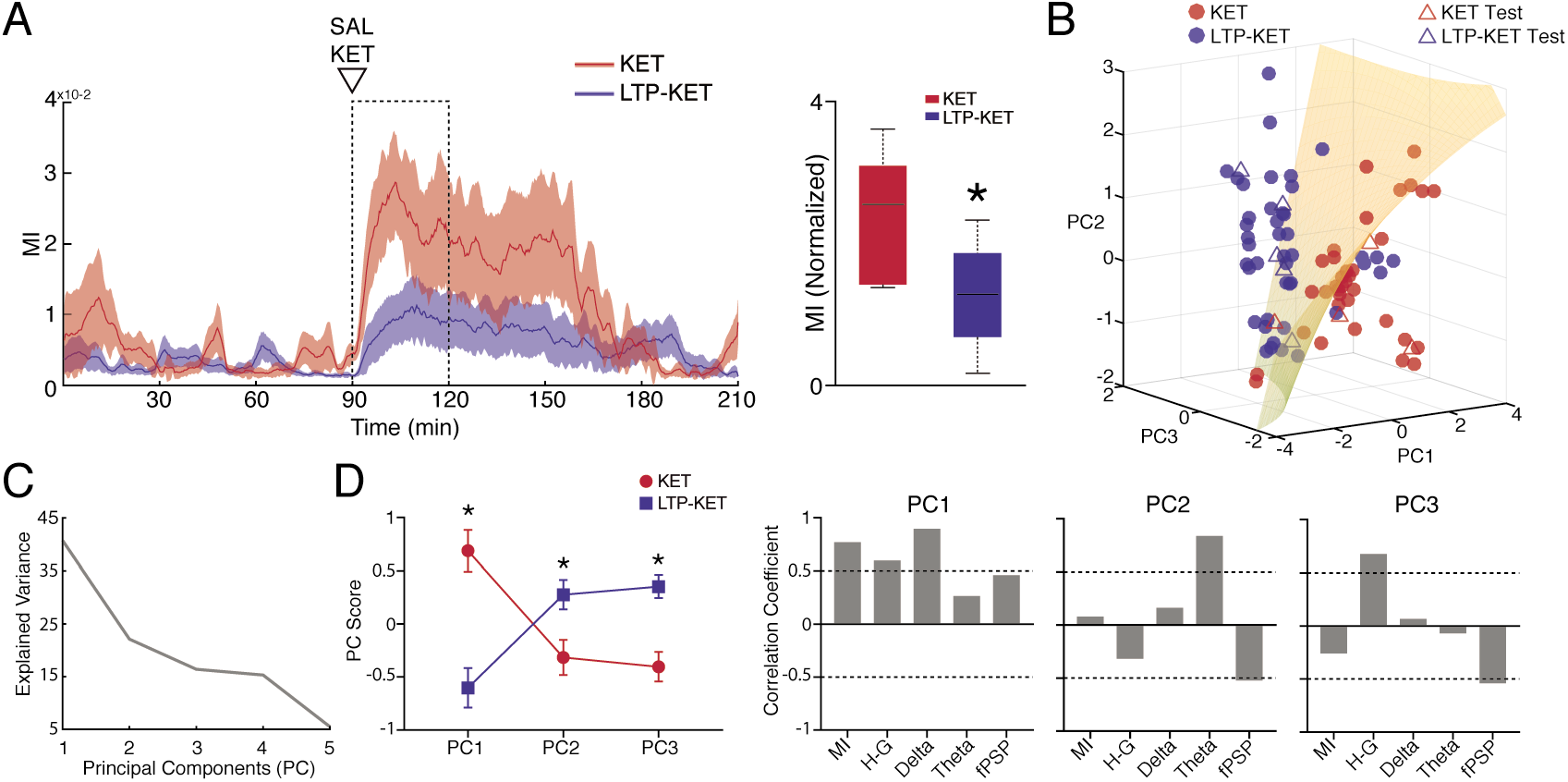
LTP induction prevents Ketamine effects. (A) LTP prevents the increase in MI state caused by ketamine. Averaged MI across the entire recording in KET (n=7) and LTP-KET (n=9) groups (left). Comparison of the baseline normalized MI in the initial 30 min after ketamine injection. There is a significant decrease in delta-high-gamma CFC when ketamine is preceded by HFS. (B) 3D plot of the Principal Component Analysis of electrophysiological features in the KET and LTP-KET groups. A quadratic discriminant function was used to classify each 5 min epochs after ketamine injection indicating separation on the training (filled circles) and test (open triangles) PFC LFPs (cross-validation accuracy: 85.56%). (C) Explained variance of the Principal Components. (D) PC score of the KET and LTP-KET epochs (left) and correlation coefficients between variables and PCs. This data indicates that KET and LTP-KET have a clear distinct electrophysiological feature. While KET epochs are described by the presence of delta, high-gamma, increased CFC and fPSP, HFS-KET epochs are characterized by the presence of theta power, elevation of high-gamma power and a reduction of fPSP. *(p<0.05).

## Discussion

In this study, we demonstrated that a prior enhancement of synaptic efficacy at hippocampal-prefrontal projections is sufficient to attenuate the disturbing effects of ketamine on oscillatory coupling and basal synaptic transmission in the cortex. Ketamine was shown to increase gamma frequency power and delta-gamma phase-amplitude coupling in the PFC and boosted HPC-PFC synaptic plasticity with PPF disruption. We also observed that LTP induction was associated with an increase of gamma power in DEA states in the PFC and no alteration in delta-gamma coupling in the PFC. Under urethane anesthesia rodents show a spontaneous alternation of brain states characterized by activated and deactivated periods^43,44^. Here, the NMDAr antagonism by S+ Ketamine (12,5 mg/Kg) changed the brain state dynamics by (1) increasing the numbers of DEA states; and (2) inducing a distinct DEA state with high gamma power and abnormal cross-frequency coupling between delta phase and high-gamma amplitude.

The induction of aberrant gamma oscillations in cortical and subcortical regions is a typical effect of acute ketamine treatment^45^. A variety of *in vivo* studies have demonstrated that non-competitive NMDAr antagonism increases the firing rate of PFC pyramidal neurons reducing their synchrony, and enhancing broadband gamma activity^33,46–49^. This broadband gamma enhancement is thought of as an aberrant and diffuse noise at the network level that can cause dysfunction in cognitive and sensory-motor integration^17,50^. In our study, gamma increase after KET injection was robust in the PFC, with subtle effects in the HPC (see also Figure S1A). Robust gamma increases have been previously reported in the HPC following NMDAr antagonism in freely moving^51,52^ and anesthetized animals^17^ as well. To our knowledge, none of these studies tested the effects of S+ ketamine, which has a higher affinity to NMDAr compared to racemic ketamine ^53^. Indeed, different NMDAr antagonists are known to produce different effects on gamma oscillatory dynamics^46,54,55^.

Interestingly, we show that in DEA states, the increase in high-gamma activity induced by ketamine is strongly coordinated by slow oscillations, rather than by a generalized increase of activity. We suggest that this rhythmic coupling is independent of enhancement in gamma activity since there was low correlation between gamma power and CFC. Post-ketamine epochs of similar gamma power to pre-ketamine showed significantly higher delta-gamma coupling. Indeed, it was previously shown that ketamine could affect CFC between theta and gamma oscillations in the hippocampus of freely moving rats^51^. In contrast, delta-gamma CFC has been described in cortico-striatal network and could reflect cortico-mesolimbic connectivity^55^, induced in frontal regions by anesthetic doses of ketamine^56^ and modulated by dopaminergic activation^57,58^. One possible mechanism by which KET produces an enhancement in CFC is by increasing extracellular levels of dopamine. Consistent with this hypothesis, it has been shown that racemic ketamine promotes dopamine release in PFC of rats^59^ and that S+ KET strongly inhibits dopamine transporter^60^. Besides, S+ KET administration is known to increase dopamine levels in the striatum of healthy subjects^61^.

Local disinhibition induced by ketamine could also underpin the sustained increase in HPC-PFC synaptic efficiency after S+ KET administration. Consistent with our findings, Blot et al. showed that under urethane anesthesia, systemic injection of MK-801 produced an increase of PFC evoked responses induced by ventral HPC stimulation^32^. In our study we observed a sustained reduction in PPF following S+ KET administration. Similarly, Kiss et al. observed a stronger but shorter reduction of HPC-PFC PPF after systemic MK-801^26^. They also noticed that changes in PPF occurred following systemic or mediodorsal thalamic nucleus microinjections, but not following injection into the PFC. This suggests that NMDAr antagonism could exert part of its effects by acting on thalamic nuclei. Given the critical role of the thalamus in integrating and regulating sensory information to the cortex, and the ability of short-term synaptic plasticity to influence information processing, it is plausible that dysfunctional sensory/cognitive processing in schizophrenia may arise from modified short-term synaptic plasticity^62–64^. Indeed, short-term synaptic plasticity alterations are a consistent finding in genetic mouse models of schizophrenia (for a review: Ruggiero et al. 2017^65^; Sigurdsson et al. 2016^66^)

Another finding of the present study was that LTP induction in the HPC-PFC projections abolished ketamine reduction of PPF. Classically, PPF arises from calcium accumulation in the presynaptic terminal due to a first stimulation pulse that results in increased neurotransmitter release in response to a second stimulus followed by a short interval^30^. However, short-term synaptic plasticity in the HPC-PFC pathway may encompass a more complex interaction, involving GABAergic interneurons terminating in the PFC pyramidal cells, since CA1 collaterals innervate both pyramidal and GABAergic neurons^67^. Electrical stimulation of CA1 induces a relevant burst activity in PFC interneurons in contrast to the few spikes elicited in pyramidal neurons^68^. This feed-forward inhibition is proposed to constrain the excitatory influence of the HPC on PFC pyramidal cells, supporting rhythmic synchronization between the hippocampus and cortical activity^68^. It is possible that PPF deficits induced by NMDAr antagonism could contribute to a reduced synchrony between HPC and PFC and LTP induction could attenuate this effect. Also, the decrease in HPC-PFC pathway PPF response is related to an increase of delta activity in the 0.5-2 Hz band^26^. Interestingly, we show that ketamine produced an increase in the number of deactivated epochs that are characterized by ~1Hz power. In contrast, LTP induction prevented this effect on brain state alternation, which could explain the reduction of KET effects on PPF.

Moreover, LTP induction attenuated the enhancement of cortical fPSP induced by ketamine injection. Blot et al. 2015 obtained a similar result in the ventral HPC-PFC projections. The authors argued that ketamine might induce plasticity in HPC-PFC synapses by mechanisms similar to LTP rather than through local synaptic disinhibition of PFC pyramidal neurons^69^. Supporting this idea, microinjection of the NMDAr competitive antagonist, AP5 abolished both MK-801 and tetanus stimulation effects on cortical fPSP^32^. In another study, a low dose of ketamine reduced the LTP magnitude induced by ventral CA1 stimulation, an effect that was prevented by clozapine administration^18^. Together, these results suggest direct competitive mechanisms: ketamine blocks active NMDAr, and LTP in the HPC-PFC pathway depends on NMDAr activation^18,69^.

Extending the effects of deep brain stimulation, we observed that cortical LTP induction increased gamma activity early after HFS (0-30 min) specifically in DEA epochs. This effect is consistent with previous reports that show gamma increase in the posterior HPC-PFC pathway following LTP, but no LTD^70^. However, we did not observe enhanced CFC following gamma increase. These findings suggest that in contrast to KET, LTP promotes less coordinated gamma oscillations during slow delta activity. Furthermore, HFS attenuated aberrant delta-high-gamma CFC during DEA epochs. As described, the efficacy of this stimulation in preventing ketamine effects could be related to an increase of AMPAr and NMDAr functions in PFC interneurons^68,71^. However, we cannot discard the contribution of other regions and modulatory systems. For instance, it is well-known that HFS delivery to the HPC causes dopamine release in the PFC^72^. Given the possibility that KET increases CFC coupling in delta range increasing dopaminergic activity, future experiments modulating dopaminergic receptors in the PFC during LTP and NMDAr antagonist administration would be elucidative.

As a demonstration that deep brain stimulation significantly changes the neural dynamics induced by KET, we were able to separate these two brain states (KET and LTP-KET) using an unsupervised algorithm solely based on the electrophysiological features of each state. It remains an open question, however, whether LTP induction would also attenuate functional and behavioral deficits induced by ketamine. Indirect supporting evidence can be obtained from reports using allosteric modulators of NMDAr or AMPAr. Application of LY451395, an AMPA positive allosteric modulator, reverts the increase in slow oscillations and PPF deficits produced by MK-801^26^. Inhibitors of glycine transporter were able to revert the psychotomimetic effects of ketamine in rodents and in healthy humans^24,73^. Nevertheless, blocking glycine reuptake in patients with schizophrenia by using bitopertin, did not improve their cognitive and negative symptoms^27,28^. Direct support, on the other hand, comes from experimental deep brain stimulation (DBS) studies. High-frequency stimulation of the ventral HPC was shown to normalize auditory evoked responses in the MAM model of schizophrenia^74^. Using the same animal model, Perez et al. 2013 showed that ventral hippocampal DBS: (1) normalized aberrant dopamine neuron activity, (2) decreased locomotor response to amphetamine, and (3) restored deficits of cognitive flexibility^75^. Furthermore, the application of DBS to other cerebral regions, such as PFC, *nucleus accumbens*^76,77^, and the medial septum^76^ showed promising results for alleviating behavioral deficits in animal models of schizophrenia. Taken together, these data suggest that HFS of limbic circuits should be further investigated as a possible treatment for drug-resistant schizophrenia.

## Conclusion

Our findings expand previous studies showing that systemic treatment with S+ KET produce complex changes in connectivity, synaptic plasticity, and oscillatory patterns in the HPC-PFC pathway *in vivo*. The prevention of most of these electrophysiological effects through LTP induction supports the idea that NMDAr antagonist effects share common mechanisms with synaptic plasticity events. Additional studies are needed to clarify the underlying molecular mechanisms of this interference induced by this form of deep brain stimulation. Additionally, our results suggest that HFS applied to the hippocampus could be a useful strategy to test the attenuation of cognitive impairments in animal models of schizophrenia. We hope these results will contribute to the development of non-pharmacological treatments aimed at preventing or mitigating cognitive deficits associated with psychiatric disorders.

## Methods

### Subjects

A total of 36 male Wistar rats weighting 300-450g were used in the experiments. Six animals were excluded based on inconsistent fPSP or mortality related with anesthesia. Rats were housed in groups of four in standard rodent cages in a controlled-temperature room (22±2 °C), on a 12h light/dark cycle (light on at 7 a.m.) with free access to food and water. All the experimental procedures were approved by the local bioethics committee (Ribeirão Preto Medical School, University of São Paulo; protocol number: 193/2009) which guidelines are in conformity with the National Institutes of Health guide for the care and use of laboratory animals (NIH Publications No. 8023, revised 1978).

### Surgical procedures and electrophysiological recordings

Animals were anesthetized with urethane (1.2-1.5 mg/Kg in NaCl 0.15 M, ip) and placed in a stereotaxic frame. After cleaning procedures the skull was exposed, and burr holes were drilled aiming the left PFC (anterior-posterior, AP: +3.0 mm; medial-lateral, ML: +0.5 mm; dorsal-ventral, DV: 3.2 mm) and HPC (CA1, AP: −4.7 mm; ML: 4 mm; DV: 2.5-2.8 mm) for recording, and HPC intermediate region (CA1, AP: −5.7 mm; ML: 4.4 mm; DV: 2.5-2.8 mm) for stimulation electrodes implant (Figure 1A). Manufactured electrodes were made of single Teflon-coated tungsten wires (60 μm, AM-Systems) for recording and two twisted wires for bipolar stimulation (~500 μm interpole distance). An epidural screw placed in the right parietal bone was used for reference and ground. Temperature (37 ± 0.5 °C) was kept constant during all the procedure by a heating pad.

HPC recording electrode positioning was adjusted by monitoring typical LFP and audio-monitor signals from the hippocampus (i.e., prominent spikes and theta oscillation). The stimulus electrode in HPC was adjusted by applying low-intensity test-pulses (square monophasic pulses, ~150 μA 200 μs, 0.05 Hz) aiming for a consistent fPSP in the PFC (i.e., the latency of first negative peak of 14-17 ms and amplitude>0.25 mV^78,79^. After electrode adjustment, an input-output curve (I/O curve; 60-500 μA) was used to establish the current intensity necessary to evoke 70% of the maximal fPSP amplitude. This current was used to apply paired monophasic pulses (same parameters as in test pulses and 80 ms of inter-pulse interval; S88; Grass Instruments) during the entire experiment. Recorded signals consisted of evoked fPSP and concomitant LFPs of PFC and HPC (Figure 1C). Signal was amplified 100x, band-pass filtered (0.3-1000 Hz; Grass), and digitized at 10 kHz (ADInstruments). LTP induction was induced by applying a high-frequency stimulation (HFS): two series of 10 trains (50 pulses at 250 Hz every 10 s) separated by 10 min.

### Experimental design

Figure 1B illustrate the experimental design. In Experiment 1 we investigated the ketamine effects on HPC-PFC connectivity. fPSPs and LFPs were monitored for 120 min after ketamine (S(+)-ketamine; 12.5 mg/Kg ip) or saline (0.9 %) injection and compared with the 90 min baseline (groups KET, n=7 and SAL, n=8, respectively). We choose S(+)-ketamine given its high affinity for NMDAr^53^ and because it reproduces the metabolic effects observed in psychotic patients^80^. In Experiment 2 we explored the hypothesis that LTP induction could prevent ketamine effects on the HPC-PFC connectivity. After a 30 min baseline, two HFS protocols were applied at 30 and 60 min. Following HFS, ketamine or saline was injected and field potentials were monitored for an additional 120 min (groups LTP-KET, n=9 and LTP-SAL, n=6, respectively). All experiments were conducted during urethane anesthesia.

### Data analysis

All data processing was performed using customized scripts in Matlab (Mathworks). The amplitude of evoked fPSP1 and fPSP2 (Figure 1C) were normalized as a percentage of baseline mean (90 and 30 min for experiments I and II, respectively). PPF was calculated as the ratio of fPSP2 and fPSP1 as an indication of short-term synaptic plasticity. All fPSP measures were averaged in blocks of 10 min. LFP signal was re-sampled to 1000 Hz and then high-pass filtered at 0.5 Hz. The whole recording data was epoched in a 20 s period following HPC electrical stimulation. Time windows of 0.5 s containing the evoked fPSP and electrical stimulation artifact (in both regions) were eliminated from all epochs.

### Brain state classification

We classified deactivated and activated periods by plotting PFC epoch values for RMS (root mean square) and the number of zero-crossing in the LFP (Figure 1D). These measures directly reflect amplitude and presence of faster rhythms in the signal (Figure 1C and D). We used a k-means algorithm (squared Euclidean distance for three groups) for an initial clustering and manually refined the classification eliminating epochs at the cluster edge. Spectrum content of each state was analyzed to confirm classification. Not classified epochs were not analyzed in this work.

### Spectral analysis

Power spectral density estimates (PSD) were calculated using Welch’s method in which Discrete Fourier Transform (FFT Matlab algorithm) is applied in overlapping windows and the periodogram is calculated for each segment individually and the magnitude squared result of the FFT is averaged. We used 3 s Hamming tapered windows, with 50 % overlap and a 2^12^ points FFT. PSDs estimates were then averaged over trials and animals. For representative spectrograms, we used a short-time Fourier Transform in the whole recording using a 60 s window with 2^16^ points FFT and 50% overlap. For statistics, power was integrated into specific frequency bands (delta: 0.5 −2 Hz, theta: 3-5 Hz, low-gamma: 30-55 Hz, and high-gamma: 65-100 Hz). Relative power was obtained by dividing PSD estimation by the integrated power over all frequencies.

Spectral coherence was estimated using Welch’s periodogram method to compute the cross-PSD of PFC and HIPO (*Pxy*) and the PSD of both region (*Pxx* and *Pyy*). The magnitude squared coherence was calculated as: *Cxy(f)* = |*Pxy(f)*|^*2*^ / *Pyy(f) Pxx(f)*. The parameters used were the same as described for the PSDs estimate. Coherograms were calculated using the same approach while using a moving window of 90 s and 50% overlap. Power and coherence values from 58 – 62 Hz (line noise contamination) were removed from the analysis. To evaluate HFS effects on spectral parameters we combined all LTP animals (LTP-SAL and LTP-KET groups) into one group (LTP group) since HFS was the only manipulation from the second electrical stimulation until drug injection.

### Phase-amplitude coupling

Cross-frequency coupling was estimated by the modulation index as described by Tort and adapted in previous work from our group^81,82^. Briefly, comodulation maps were constructed applying Hilbert transform to the signal filtered in bins of 0.5 Hz from 1 to 20 Hz on steps of 1 Hz for the phase modulating signal, while the amplitude modulated signal was filtered in bins of 1 Hz from 10 to 120 Hz on steps of 5 Hz. Shannon entropy of the distribution of mean amplitudes per phase (divided into 18 bins) in each frequency bin was calculated to obtain the cross-frequency modulation index (MI) for each period. The MI between delta oscillations (1-2 Hz) and high gamma band (65-100 Hz) was calculated for comparisons.

### Directionality analysis

We inferred directionality by cross-correlation and Granger causality. Cross-correlation measures the similarity of two time series by performing the sliding dot product between the signals. We calculated the peak lag of the cross-correlation of delta and theta filtered signals for deactivated and activated epochs, respectively. Wiener-Granger causality spectra were performed using the MVGC toolbox developed by Barnett and Seth^83^, which is freely available online (http://users.sussex.ac.uk/~lionelb/MVGC/). Such algorithm uses vector auto-regressive models to estimate prediction of a time series A based on another time series B comparing with the prediction obtained by using the past values of time series A alone. We used pairs of HPC and PFC LFP separating for DEA or ACT epochs. Initially, raw LFP were decimated to 200 Hz, and the model order was estimated by Akaike Information Criterion for each animal separately (model order range: 36-38). We fixed the model order of 40, which gave an adequate frequency resolution for the slow oscillations that predominate on our signals with reasonable computation cost. For statistical significance, we calculated the 95 % confidence interval (CI) from the empirical null distribution of the frequency-domain Granger estimates, based on randomly permuting one LFP in bins the size of the model order. For directionality comparison (HPC→PFC vs. PFC→HPC; in DEA or ACT) we used a Bonferroni-corrected paired *t* test for the peak frequency in the delta (0.5 – 2 Hz) or theta (3 – 5 Hz) bands.

### Principal Component and Discriminant Analysis

Principal component analysis, using singular-value decomposition (*pca* Matlab function), was used for dimensionality reduction in order to find patterns of variance among multivariate data. Variables analyzed were MI, high-gamma, delta and theta power and fPSP amplitude. We used these features since they were associated with ketamine effects (Figure 3). LFP data were normalized by the mean value of baseline DEA epochs, while for fPSP we used the normalized mean value for the 10 min before drug injection (all epochs regardless of state classification to account for LTP induction). The normalized values were extracted in 5 min epochs from 10-40 min after drug injection (KET and LTP-KET groups) in order to capture stable effects of ketamine. Data was z-scored for each variable. Following PCA analysis, data was projected against principal components (PCs) and the mean score was compared between conditions (KET vs. LTP-KET). Correlation coefficients between original variables and scaled components were obtained by multiplying eigenvectors by the square root of the eigenvalues. For interpretation, we used correlation coefficients value >0.5^84^. A discriminant analysis classifier was used based on the first three principal components extracted (explained variance of 79.21%). A quadratic function fit was heuristically determined and a 50 fold cross-validation was performed with 81 epochs for training data and 9 epochs for data test. Crossvalidation of the quadratic discriminant model using PC1-PC3 resulted in 85.56% accuracy, which was better than the 83.33% cross-validation accuracy of the discriminant fit using all the 5 dimensions of the original data.

### Statistics

Normal distribution was evaluated in all data sets using Kolmogorov-Smirnov test. fPSP data were analyzed by two-way ANOVA with repeated measures and Bonferroni post-hoc test to compare treatment over time. For LFP (power and CFC) we used paired *t* tests for within-group comparisons, and unpaired *t* tests for between-group comparison or Wilcoxon matched-pairs signed rank and Mann-Whitney test, respectively, as non-parametric equivalents. For comparison of more than two conditions we used one-way ANOVA with tukey-kramer post-hoc or Kruskal-Wallis test with Dunn’s post-hoc for non-Gaussian distribution. Significance of spectral coherence and MI were estimated calculating a CI for a surrogate data using a bootstrapped shuffled data (8000 iterations). Probability density function after drug injection was estimated calculating the presence of DEA classified epochs for all the animals in Saline or ketamine groups. Pearson’s correlation coefficient was calculated to investigate linear dependency between CFC and LFP power. Data are expressed as the mean ± standard error of the mean (SEM) for bar and line plots, and for Box-plot data are expressed as 1^st^ quartiles, medians, and 3^rd^ quartiles, with whiskers representing minimum and maximum values. The significance level was set to 0.05.

### Histology

To confirm electrode positioning we performed an electrolytic lesion (1mA, 1s) at the end of the trial. After an additional dose of anesthesia animals were decapitated and had their brains removed and placed in solutions for fixation (10% formaldehyde in phosphate-buffered saline, PBS) and cryoprotection (20% sucrose in PBS). Coronal sections (30 μm) stained with cresyl violet were evaluated through a bright-field microscope (Figure 1A).

### Data availability

The datasets generated during and/or analyzed during the current study are available from the corresponding author on reasonable request.

## Acknowledgements

This work was funded by the São Paulo Research Foundation, FAPESP (2014/18211-0 for C.L.A., 2018/02303-4 for R.N.R., and 2016/17882-4 for J.P.L.), the National Council for Scientific and Technological Development, CNPq (142451/2014-2 for M.T.R., 481629/2011-4 for R.N.R.P. and 466995/2014-8 for J.P.L.). We thank Antônio Renato M. Silva and Renata C. Scandiuzzi for their assistance. We thank Danilo Benette Marques, Eliezyer Fermino de Oliveira and Stephanie Rogers for the valuable comments on this manuscript.

## Author contributions statement

C.L.A., R.N.R.P. and J.P.L. conceived the experiments, C.L.A., M.T.R, R.N.R., I.M.E. and J.E.P.S conducted the experiments, M.T.R. prepared the figures, R.N.R., C.L.A. and R.N.R.P. analyzed the data, R.N.R., C.L.A. and M.T.R. wrote the manuscript. All authors reviewed the manuscript.

## Supplementary figures

**Figure S1 –.**
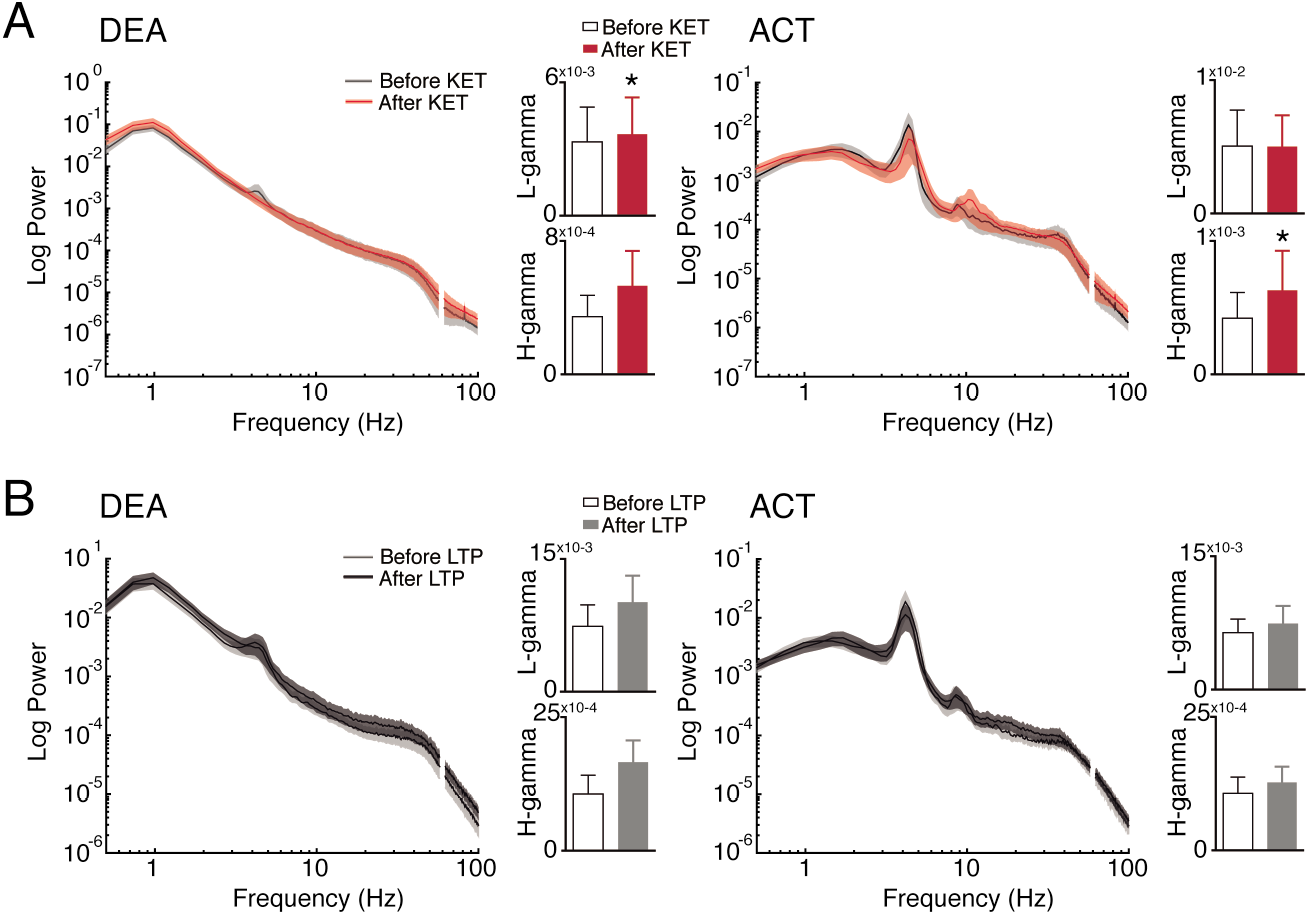
KET and HFS effects on HPC power spectral density. (A) Ketamine produced a subtle low-gamma increase in DEA (left, Wilcoxon test, n=7, *p*=0.0469) and high-gamma (right, Wilcoxon test, n=7, *p*=0.0312) in ACT states (right). (B) LTP induction did not produce a gamma increase in DEA (left) or ACT states (right). Line plots representing average PSDs in the HPC before and after ketamine injection. Bar plots representing low and high-gamma frequency bands before and after ketamine administration. *(p<0.05).

**Figure S2 –.**
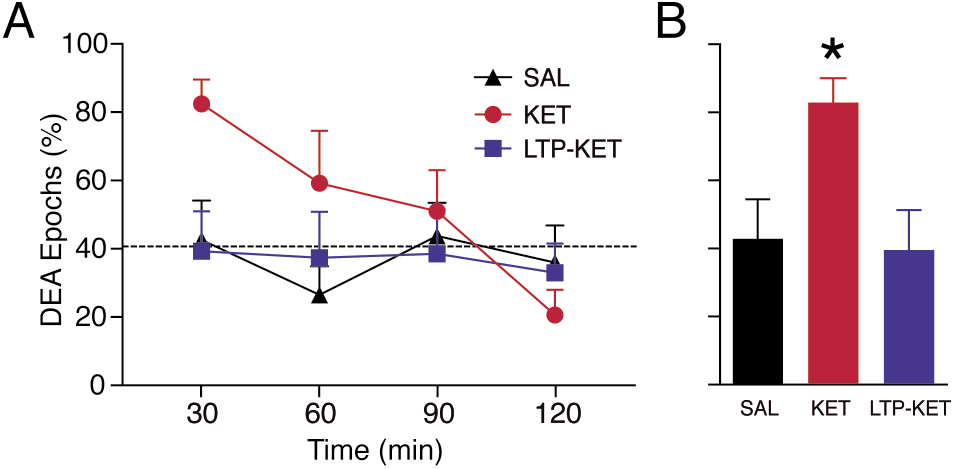
Ketamine administration effects on the brain state alternation under urethane anesthesia. (A) Time-course of the average percentage of DEA epochs percentage after KET or Sal administration (left). The dashed line represents the chance of obtaining a DEA epoch in a 30 min period calculated by permuting all epochs from all groups 40,000 times. (B) Average percentage of DEA epochs in the initial 30 min after drug administration. Ketamine produces an acute (30 min) increase in brain states epochs classified as DEA. One-way ANOVA: F_(2,21)_=4.64, *p*=0.0214, Tukey post-hoc test *p*=0.049 for KET vs. Sal, and *p*=0.0271 for KET vs. LTP-KET. Interestingly, LTP induction prevented KET increase in DEA epochs. *(p<0.05).

## References

1. Godsil, B. P., Kiss, J. P., Spedding, M. & Jay, T. M. The hippocampal–prefrontal pathway: The weak link in psychiatric disorders? Eur. Neuropsychopharmacol. 23, 1165–1181 (2013).

2. Laroche, S., Davis, S. & Jay, T. M. Plasticity at hippocampal to prefrontal cortex synapses: Dual roles in working memory and consolidation. Hippocampus 10, 438–446 (2000).

3. Bähner, F. & Meyer-lindenberg, A. Hippocampal – prefrontal connectivity as a translational phenotype for schizophrenia. Eur. Neuropsychopharmacol. 27, 93–106 (2017).

4. Cohen, S. M., Tsien, R. W., Goff, D. C. & Halassa, M. M. The impact of NMDA receptor hypofunction on GABAergic neurons in the pathophysiology of schizophrenia. Schizophr. Res. 167, 98–107 (2015).

5. Schneider, M. et al. Altered DLPFC–hippocampus connectivity during working memory: Independent replication and disorder specificity of a putative genetic risk phenotype for schizophrenia. Schizophr. Bull. 43, 1114–1122 (2017).

6. Meyer-lindenberg, A. S. et al. Regionally specific disturbance of dorsolateral prefrontal–hippocampal functional connectivity in schizophrenia. Arch. Gen. Psychiatry 62, 379–386 (2005).

7. Whitfield-gabrieli, S. et al. Hyperactivity and hyperconnectivity of the default network in schizophrenia and in first-degree relatives of persons with schizophrenia. Proc. Natl. Acad. Sci. 106, 1279–1284 (2009).

8. Pilowsky, L. S. et al. First in vivo evidence of an NMDA receptor deficit in medication-free schizophrenic patients. Mol. Psychiatry 11, 118–119 (2006).

9. Catts, V. S., Ling, Y., Shannon, C., Weickert, T. W. & Catts, S. V. A quantitative review of the postmortem evidence for decreased cortical N-methyl-d-aspartate receptor expression levels in schizophrenia: How can we link molecular abnormalities to mismatch negativity deficits? Biol. Psychol. 116, 57–67 (2016).

10. Grimm, O. et al. Acute ketamine challenge increases resting state prefrontal-hippocampal connectivity in both humans and rats. Psychopharmacology (Berl). 232, 4231–4241 (2015).

11. Obi-Nagata, K., Temma, Y. & Hayashi-Takagi, A. Synaptic functions and their disruption in schizophrenia: From clinical evidence to synaptic optogenetics in an animal model. Proc. Japan Acad. Ser. B Phys. Biol. Sci. 95, 179–197 (2019).

12. Penadés, R., Franck, N., González-Vallespí, L. & Dekerle, M. Reviews on Biomarker Studies in Psychiatric and Neurodegenerative Disorders. Adv. Exp. Med. Biol. 1118, 118–134 (2019).

13. Haaf, M., Leicht, G., Curic, S. & Mulert, C. Glutamatergic deficits in schizophrenia – Biomarkers and pharmacological interventions within the ketamine model. Curr. Pharm. Biotechnol. 19, 293–307 (2018).

14. Bondi, C., Matthews, M. & Moghaddam, B. Glutamatergic animal models of schizophrenia. Curr. Pharm. Des. 18, 1593–1604 (2012).

15. Lee, G. & Zhou, Y. NMDAR hypofunction animal models of schizophrenia. Front. Mol. Neurosci. 12, 1–26 (2019).

16. Kamiyama, H. et al. Mechanisms underlying ketamine-induced synaptic depression in rat hippocampus-medial prefrontal cortex pathway. Neuroscience 177, 159–169 (2011).

17. Hakami, T. et al. NMDA receptor hypofunction leads to generalized and persistent aberrant gamma oscillations independent of hyperlocomotion and the state of consciousness. PLoS One 4, (2009).

18. Rame, M. et al. Clozapine counteracts a ketamine-induced depression of hippocampal-prefrontal neuroplasticity and alters signaling pathway phosphorylation. PLoS One 12, 1–20 (2017).

19. Moghaddam, B. & Javitt, D. From revolution to evolution: The glutamate hypothesis of schizophrenia and its implication for treatment. Neuropsychopharmacology 37, 4–15 (2012).

20. Ranganathan, M. et al. Attenuation of ketamine-induced impairment in verbal learning and memory in healthy volunteers by the AMPA receptor potentiator PF-04958242. Mol. Psychiatry 22, 1633–1640 (2017).

21. Goff, D. C. et al. A placebo-controlled add-on trial of the ampakine, CX516, for cognitive deficits in schizophrenia. Neuropsychopharmacology 33, 465–472 (2008).

22. Roberts, B. M. et al. Prevention of ketamine-induced working memory impairments by AMPA potentiators in a nonhuman primate model of cognitive dysfunction. Behav. Brain Res. 212, 41–48 (2010).

23. Chen, L., Muhlhauser, M. & Yang, C. R. Glycine tranporter-1 blockade potentiates NMDA-mediated responses in rat prefrontal cortical neurons in vitro and in vivo. J. Neurophysiol. 89, 691–703 (2003).

24. Javitt, D. C. Glutamate as a therapeutic target in psychiatric disorders. Mol. Psychiatry 9, 984–997 (2004).

25. Matsumoto, M. et al. Characterization of clozapine-induced changes in synaptic plasticity in the hippocampal – mPFC pathway of anesthetized rats. Brain Res. 1195, 50–55 (2008).

26. Kiss, T., Hoffmann, W. E. & Hajós, M. Delta oscillation and short-term plasticity in the rat medial prefrontal cortex: modelling NMDA hypofunction of schizophrenia. Int. J. Neuropsychopharmacol. 14, 29–42 (2011).

27. Bugarski-kirola, D. et al. Bitopertin in negative symptoms of schizophrenia — results from the phase III flashlyte and daylyte studies. Biol. Psychiatry 82, 8–16 (2017).

28. Kantrowitz, J. T. et al. Neurophysiological effects of bitopertin in schizophrenia. J. Clin. Psychopharmacol. 37, 447–451 (2017).

29. Lüscher, C. & Malenka, R. C. NMDA receptor-dependent long-term potentiation and longterm depression (LTP/LTD). Clod Spring Harb. Perspect. Biol. 4, 1–16 (2012).

30. Citri, A. & Malenka, R. C. Synaptic plasticity: Multiple forms, functions, and mechanisms. Neuropsychopharmacology 33, 18–41 (2008).

31. Jodo, E. The role of the hippocampo-prefrontal cortex system in phencyclidine-induced psychosis: A model for schizophrenia. J. Physiol. - Paris 107, 434–440 (2013).

32. Blot, K. et al. Modulation of hippocampus–prefrontal cortex synaptic transmission and disruption of executive cognitive functions by MK-801. Cereb. Cortex 25, 1348–1361 (2015).

33. Molina, L. A., Skelin, I. & Gruber, A. J. Acute NMDA receptor antagonism disrupts synchronization of action potential firing in rat prefrontal cortex. PLoS One 9, 1–10 (2014).

34. Gean, P., Chang, F.-C., Huang, C.-C., Lin, J.-H. & Way, L.-J. Long-term enhancement of EPSP and NMDA receptor-mediated synaptic transmission in the amygdala. Brain Res. Bull. 31, 7–11 (1993).

35. Maren, S., Tocco, G., Standley, S., Baudry, M. & Thompson, R. F. Postsynaptic factors in the expression of long-term potentiation (LTP): Increased glutamate receptor binding following LTP induction in vivo. Proc. Natl. Acad. Sci. U. S. A. 90, 9654–9658 (1993).

36. Grosshans, D. R., Clayton, D. A., Coultrap, S. J. & Browning, M. D. LTP leads to rapid surface expression of NMDA but not AMPA receptors in adult rat CA1. Nat. Neurosci. 5, 27–33 (2002).

37. Pagliardini, S., Gosgnach, S. & Dickson, C. T. Spontaneous sleep-like brain state alternations and breathing characteristics in urethane anesthetized mice. PLoS One 8, 1–10 (2013).

38. Pagliardini, S., Funk, G. D. & Dickson, C. T. Breathing and brain state: Urethane anesthesia as a model for natural sleep. Respir. Physiol. Neurobiol. 188, 324–332 (2013).

39. Sapkota, K. et al. GluN2D N-methyl-D-aspartate receptor subunit contribution to the stimulation of brain activity and gamma oscillations by ketamine: Implications for schizophrenia. J. Pharmacol. Exp. Ther. 356, 702–711 (2016).

40. Lopes-Aguiar, C. et al. Muscarinic acetylcholine neurotransmission enhances the late-phase of long-term potentiation in the hippocampal-prefrontal cortx pathway of rats in vivo: a possible involvement of monoaminergic systems. Neuroscience 153, 1309–1319 (2008).

41. Lopes-aguiar, C., Bueno-junior, L. S., Naime, R., Romcy-pereira, R. N. & Pereira, J. Neuropharmacology NMDA receptor blockade impairs the muscarinic conversion of subthreshold transient depression into long-lasting LTD in the hippocampus e prefrontal cortex pathway in vivo: Correlation with gamma oscillations. Neuropharmacology 65, 143–155 (2013).

42. Esteves, I. M. et al. Chronic nicotine attenuates behavioral and synaptic plasticity impairments in a streptozotocin in a model of Alzheimer’s disease. Neuroscience 353, 87–97 (2017).

43. Kiss, T., Feng, J., Hoffmann, W. E., Shaffer, C. L. & Hajós, M. Rhythmic theta and delta activity of cortical and hippocampal neuronal networks in genetically or pharmacologically induced N-methyl-D-Aspartete receptor hypofunction under urethane anesthesia. Neuroscience 237, 255–267 (2013).

44. Clement, E. A. et al. Cyclic and sleep-like spontaneous alternations of brain state under urethane anaesthesia. PLoS One 3, 1–15 (2008).

45. Kocsis, B., Brown, R. E., Mccarley, R. W. & Hajos, M. Impact of ketamine on neuronal network dynamics: Translational modeling of schizophrenia-relevant deficits. CNS Neurosci. Ther. 19, 437–447 (2013).

46. Pinault, D. N-methyl D-aspartate receptor antagonists ketamine and MK-801 induce wake-related aberrant gama oscillations in the rat neocortex. Biol. Psychiatry 63, 730–735 (2008).

47. Anver, H., Ward, P. D., Magony, A. & Vreugdenhil, M. NMDA receptor hypofunction phase couples independent gamma-oscillations in the rat visual cortex. Neuropsychopharmacology 36, 519–528 (2010).

48. Wood, J., Kim, Y. & Moghaddam, B. Disruption of prefrontal cortex large scale neuronal activity by different classes of psychotomimetic drugs. J. Neurosci. 32, 3022–3031 (2012).

49. Carlé, M. et al. A critical role for NMDA receptors in parvalbumin interneurons for gamma rhythm induction and behavior. Mol. Psychiatry 17, 537–548 (2011).

50. Sohal, V. S. & Rubenstein, J. L. R. Excitation-inhibition balance as a framework for investigating mechanisms in neuropsychiatric disorders. Mol. Psychiatry 24, 1248–1257 (2019).

51. Caixeta, F. V, Cornélio, A. M., Scheffer-teixeira, R., Ribeiro, S. & Tort, A. B. L. Ketamine alters oscilatory coupling in the hippocampus. Sci. Rep. 3, 1–10 (2013).

52. Ye, T. et al. Ten-hour exposure to low-dose ketamine enhances corticostriatal crossfrequency coupling and hippocampal broad-band gamma oscillations. Front. Neural Circuits 12, 1–16 (2018).

53. Oye, I., Paulsen, O. & Maurset, A. Effects of ketamine on sensory perception: Evidence for a role of N-methyl-D-aspartate receptors. J. Pharmacol. Exp. Ther. 260, 1209–1213 (1992).

54. Phillips, K. G. et al. Differential effects of NMDA antagonists on high frequency and gamma EEG oscillations in a neurodevelopmental model of schizophrenia. Neuropharmacology 62, 1359–1370 (2012).

55. Pittman-polletta, B., Hu, K. & Kocsis, B. Subunit-specific NMDAR antagonism dissociates schizophrenia subtype-relevant oscillopathies associated with frontal hypofunction and hippocampal hyperfunction. Sci. Rep. 8, 1–14 (2018).

56. Pal, D. et al. Propofol, sevoflurane, and ketamine induce a reversible increase in delta-gamma and theta-gamma phase-amplitude coupling in frontal cortex of rat. Front. Syst. Neurosci. 11, 1–13 (2017).

57. Andino-pavlovsky, V. et al. Dopamine modulates delta-gamma phase-amplitude coupling in the prefrontal cortex of behaving rats. Front. Neural Circuits 11, 1–13 (2017).

58. López-azcárate, J. et al. Delta-mediated cross-frequency coupling organizes oscillatory activity across the rat cortico-basal ganglia network. Front. Neural Circuits 7, 1–16 (2013).

59. Lorrain, D. S., Baccei, C. S., Bristow, L. J., Anderson, J. J. & Varney, M. A. Effects of ketamine and N-methyl-D-Aspartete on glutamete and dopamine release in the rat prefrontal cortex: modulation by a group II slective metabotropic glutamate receptor agonist LY379268. Neuroscience 117, 697–706 (2003).

60. Nishimura, M. & Sato, K. Ketamine stereoselectively inhibits rat dopamine transporter. Neurosci. Lett. 274, 131–134 (1999).

61. Pagliardini, S., Greer, J. J., Funk, G. D. & Dickson, C. T. State-dependent modulation of breathing in urethane-anesthetized rats. J. Neurosci. 32, 11259–11270 (2012).

62. Crabtree, G. W. & Gogos, J. A. Synaptic plasticity, neural circuits, and the emerging role of altered short-term information processing in schizophrenia. Front. Synaptic Neurosci. 6, 1–27 (2014).

63. Abbott, L. F. & Regehr, W. G. Synaptic computation. Nature 431, 796–803 (2004).

64. Mongillo, G., Barak, O. & Tsodyks, M. Synaptic theory of working memory. Science (80-.). 319, 1543–1547 (2008).

65. Ruggiero, R. N. et al. Cannabinoids and vanilloids in schizophrenia: Neurophysiological evidence and directions for basic research. Front. Pharmacol. 8, 1–27 (2017).

66. Sigurdsson, T. Neural circuit dysfunction in schizophrenia: Insights from animal models. Neuroscience 321, 42–65 (2016).

67. Gabbott, P., Headlam, A. & Busby, S. Morphological evidence that CA1 hippocampal afferents monosynaptically innervate PV-containing neurons and NADPH-diaphorase reactive cells in the medial prefrontal cortex (Areas 25 / 32) of the rat. Brain Res. 946, 314–322 (2002).

68. Tierney, P. L., Dégenètais, E., Thierry, A., Glowinski, J. & Gioanni, Y. Influence of the hippocampus on interneurons of the rat prefrontal cortex. Eur. J. Neurosci. 20, 514–524 (2004).

69. Jay, T. M., Burette, F. & Laroche, S. NMDA receptor-dependent long-term potentiation in the hippocampal afferent fibre system to the prefrontal cortex in the rat. Eur. J. Neurosci. 7, 247–250 (1995).

70. Izaki, Y. & Akema, T. Gamma-band power elevation of prefrontal local field potential after posterior dorsal hippocampus – prefrontal long-term potentiation induction in anesthetized rats. Exp. Brain Res. 184, 249–253 (2008).

71. Homayoun, H. & Moghaddam, B. NMDA receptor hypofunction produces opposite effects on prefrontal cortex interneurons and pyramidal neurons. J. Neurosci. 27, 11496–11500 (2007).

72. Jay, T. M. et al. Plasticity at hippocampal to prefrontal cortex synapses is impaired by loss of dopamine and stress: Importance for psychiatric diseases. Neurotox. Res. 6, 233–244 (2004).

73. Yang, S. Y., Hong, C. J., Huang, Y. H. & Tsai, S. J. The effects of glycine transporter I inhibitor, N-methylglycine (sarcosine), on ketamine-induced alterations in sensorimotor gating and regional brain c-Fos expression in rats. Neurosci. Lett. 469, 127–130 (2010).

74. Ewing, S. G. & Grace, A. A. Deep brain stimulation of the ventral hippocampus restores deficits in processing of auditory evoked potentials in a rodent developmental disruption model of schizophrenia. Schizophr. Res. 143, 377–383 (2013).

75. Perez, S. M., Shah, A., Asher, A. & Lodge, D. J. Hippocampal deep brain stimulation reverses physiological and behavioural deficits in a rodent model of schizophrenia. Int. J. Neuropsychopharmacol. 16, 1331–1339 (2013).

76. Bikovsky, L. et al. Deep brain stimulation improves behavior and modulates neural circuits in a rodent model of schizophrenia. Exp. Neurol. 283, 142–150 (2016).

77. Ma, J. & Leung, L. S. Deep brain stimulation of the medial septum or nucleus accumbens alleviates psychosis-relevant behavior in ketamine-treated rats. Behav. Brain Res. 266, 174–182 (2014).

78. Laroche, S., Jay, T. M. & Thierry, A. Long-term potentiation in the prefrontal cortex following stimulation of the hippocampal CA1/subicular region. Neurosci. Lett. 114, 184–190 (1990).

79. Jay, T. M., Burette, F. & Laroche, S. Plasticity of the hippocampal-prefrontal synapses. J. Physiol. - Paris 90, 361–366 (1996).

80. Vollenweider, F. X., Leenders, K. L., Hell, D. & Angst, J. Differential psychopathology and patterns of cerebral glucose utilisation produced by (S)- and (R)-ketamine in healthy volunteers using positron emission tomography (PET). Eur. Neuropharmacol. 7, 25–38 (1997).

81. Ruggiero, R. N. et al. Lithium modulates the muscarinic facilitation of synaptic plasticity and theta-gamma coupling in the hippocampal-prefrontal pathway. Exp. Neurol. 304, 90–101 (2018).

82. Tort, A. B. L., Komorowski, R., Eichenbaum, H. & Kopell, N. Measuring phase-amplitude coupling between neuronal oscillations of different frequencies. J. Neurophysiol. 104, 1195–1210 (2010).

83. Barnett, L. & Seth, A. K. The MVGC multivariate Granger causality toolbox: A new approach to Granger-causal inference. J. Neurosci. Methods 223, 50–68 (2014).

84. Budaev, S. V. Using principal components and factor analysis in animal behaviour research: Caveats and guidelines. Ethology 116, 472–480 (2010).

